# Genomic Diversity and Climate Adaptation in Brachypodium

**DOI:** 10.1101/015495

**Authors:** Pip Wilson, Jared Streich, Justin Borevitz

## Abstract

The Brachypodium genus contains the model grasses *B. distachyon*, *B. stacei* and *B. hybridum*, that are useful for molecular and physiological studies relevant to grain, pasture and bioenergy crops, as well as ecology. In this chapter we discuss the natural variation in climate/geography, genotypic and phenotypic diversity that exists within these species. We describe utilisation of this diversity via two methods, Genome Wide Association Studies and Landscape Genomics, to examine the interaction between specific genetic variants, phenotype, and environment. The aim is to identify adaptive loci that control specific traits in specific environments and understand the contribution of background polygenetic variation shaped by demographic processes. With recent developments in high throughput phenotyping, cheaper genotyping by sequencing and higher spatial/temporal resolution of climate data, these approaches can exploit the diversity of the Brachypodium. Experiments using this toolkit will reveal alleles, genes and pathways underlying agriculturally important and environmentally sensitive traits for use in grass breeding.

## Introduction

Brachypodium is part of the Poaceae plant family, one of the five largest. Poaceae are one of two plant families found on all seven continents and grow in most habitats from alpine, to freshwater, and marine environments (Groombridge and Jenkins 2002; Brummitt and Cheek 2007). The Pooideae subfamily of the Poaceae are also widespread, especially in temperate zones, and many genera extend to cold climates in mountainous regions or high latitudes. Similarly to other Pooideae genera, the Brachypodium genus also has a wide distribution in temperate regions, but is less common in cold climates and more abundant in hot arid regions (Hartley 1973). Brachypodium native range is the Mediterranean region with accessions commonly found through southern Europe, North Africa and Eurasia. However, worldwide herbarium records show Brachypodium is now present on all six continents except Antarctica (Garvin et al. 2008). This wide geographic range and climate tolerance in both the native and introduced range, as well as a key evolutionary position, mean Brachypodium is a rich resource of allelic diversity.

### 1 Diversity in Natural Populations of Brachypodium

#### Climate and Geographic Diversity

A large collection of thousands of Brachypodium accessions are now available. They come from a range of seasonal temperature profiles that span from alpine to hot arid deserts (Figure 1). The majority of accessions have been collected from temperate grasslands with hot dry summers and wet winters (Mediterranean). These include accessions from southern Europe and South-Western USA. Many accessions come from temperate regions that also get summer rainfall, such as France and South-Eastern Australia. A few accessions come from regions with colder climates, such as mountainous regions in Turkey and on the Spanish/French border. Lastly, several accessions come from arid regions in the Middle East. These different climate types favour the development of different life history strategies allowing optimization of reproductive success, as well as abiotic stress tolerance mechanisms to deal with heat, cold and drought.

**Fig. 1.**
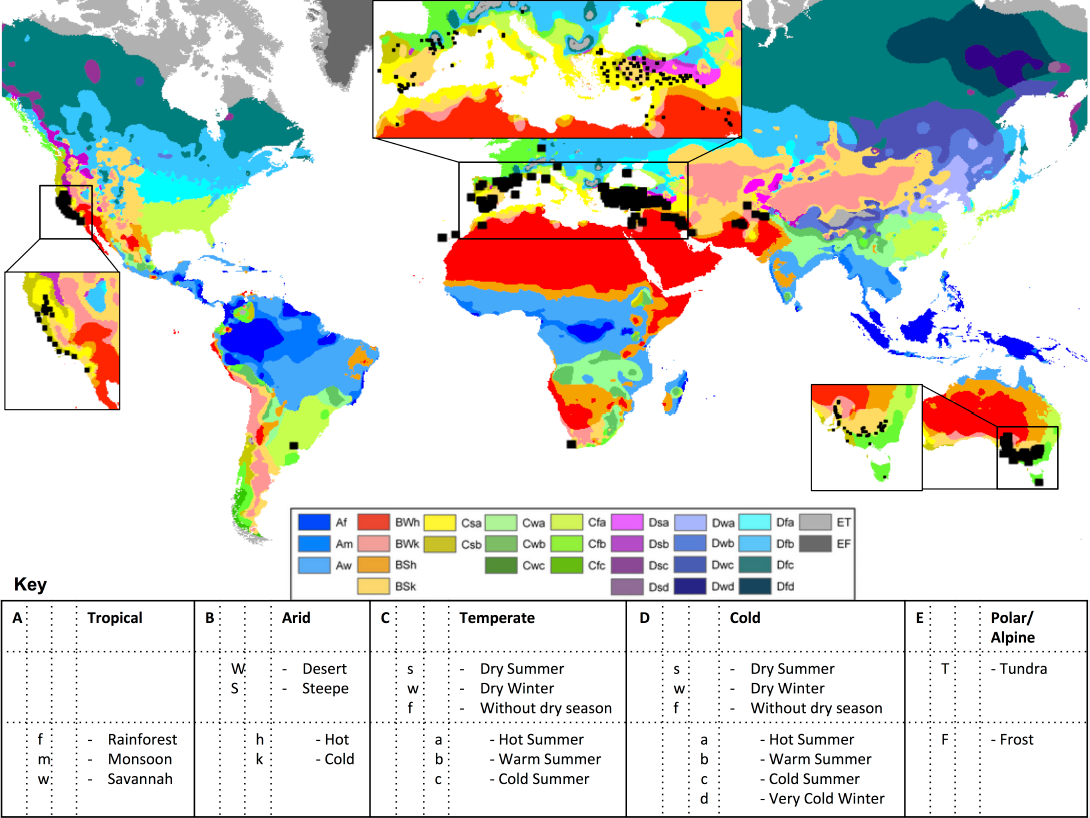
Geographic and climatic distribution of Brachypodium accessions. Distribution of the 350 populations, >1,400 accessions, of Brachypodium overlaid on the Köppen-Geiger climatic regions (Peel et al. 2007). Many other private collections exist.

The geographic distribution of Brachypodium is dependent on climate and the history of its dispersal. Brachypodium species have had a long association with human civilisation. There is evidence of use of Brachypodium species as food in Paleolithic societies in the region of Italy as long as 30 000 years ago (Revedin et al. 2010). This association may have aided its dispersal through Europe and Eurasia. In more modern times, Brachypodium co-localisation with cereal crops may have aided its dispersal through joint harvest and planting, especially from its native range in the Mediterranean to the “New World” of the Americas and Australia.

#### Genomic Diversity

There is a large amount of genomic diversity in the *Brachypodium distachyon* complex. Firstly there are three species now described. *B. distachyon* (2n=10) and *B. stacei* (2n=20) share common ancestry at large fractions of their genome and *B. hybridum* (2n=30) is an allopolyploid of *B. stacei* and *B. distachyon*. Secondly there is major population genomic structure within each species. Genotyping By Sequencing (GBS) is an efficient way to characterize population and phylogenomic variation within and between populations and species in a complex. It uses restriction digestion, barcoded adaptor ligation, pooling, second generation sequencing, and de-multiplexing to identify SNPs and call genotypes at >10,000s loci (Elshire et al. 2011). Multiplexing reduces the individual sample cost of sequencing allowing many more accessions to be sequenced, up to 384 in a single sequencing lane.

Preliminary results are showing that *B. distachyon* can be divided into two subspecies, those accessions that are similar to the Bd21 accessions, denoted as subspecies A, and those that are similar to the Bd1-1 accession, denoted as subspecies B. While the two subspecies can be intercrossed in the laboratory, the barriers seem to be maintained in nature. Within each of these subspecies there is further geographic subdivision in genetic diversity between Eastern and Western Europe (Figure 2).

**Fig. 2.**
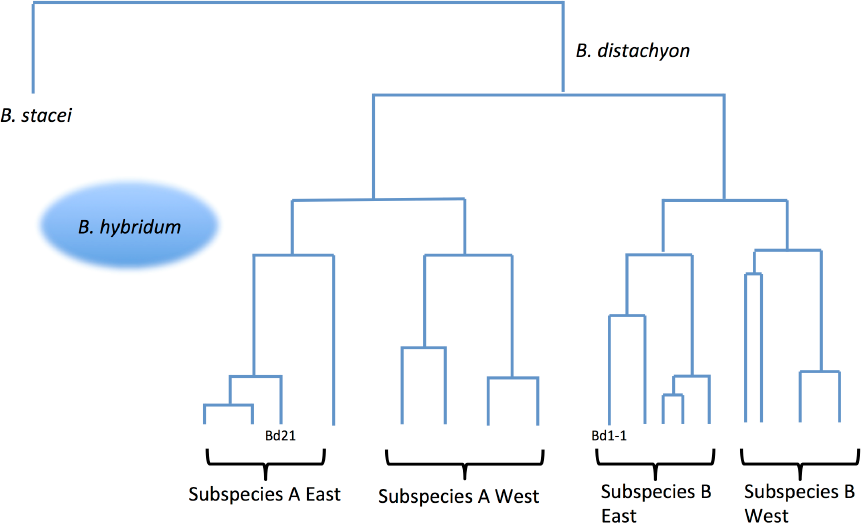
Subspecies and population genomic subdivision with Brachypodium. The first division in *B. distachyon* is that between the A and the B subspecies. Second is the geographic split between Eastern and Western Europe. This implies independent re-colonization of Europe, for each subspecies via both the east and west routes after glacial retreat.

This East-West division is most likely due to isolation by distance. Arabidopsis also shows a weak east west divide due to recolonisation of Europe, after the last ice age from refugia on the Iberian Peninsula and Western Asia (Sharbel et al. 2000). Brachypodium has a lot of parallels with Arabidopsis including its highly selfing nature, similar native range of Europe and Eurasia, similar habitats in disturbed sites and similar annual life strategy. The pattern of range retraction and subsequent recolonisation likely explains the genetic differentiation among Eastern and Western Europe genotypes seen in *Brachypodium distachyon, Arabidopsis thaliana*, and many other species. The presence of this pattern in both the A and B subspecies indicates parallel re-colonisations of subspecies that diverged before the last ice age.

Though currently under represented in germplasm collections, the evolution that created *B. stacei* appears to have one prominent haplotype, and many rare haplotypes (Catalan et al. 2012). A substantial portion of the *B. distachyon* genome is shared with *B. stacei* across many chromosome segments. *B. hybridum* are allotetraploids, resulting from a recent hybridization events fusing *B. distachyon* and *B. stacei* (Betekhtin et al. 2014). This polyploidisation with *B. stacei* may have occurred independently with both A and B subspecies of *B. distachyon* (further discussed in Catalan et al. 2012). All three Brachypodium species are morphologically similar to each other, but express different traits. *B. stacei* and *B. hybridum* have higher seed yield, more biomass and quicker growth rate than *B. distachyon* (Catalan et al. 2012; Vogel et al. 2009). They also have different ranges with the majority of accessions found in North America and Australia being *B. hybridum* while *B. stacei* has been found throughout Mediterranean regions. These species provide ideal models for studies of allopolyploidy with parallels particular to wheat, *Triticum aestivum*.

#### Adaptive Phenotypic Variation

The genetic makeup, and resulting phenotypic expression, of a given accession is affected by the genotypic and climatic history of its ancestors as this determines the suite of alleles present in its genome. Past and present selective pressures, isolation by distance, and the prevalence of intercrossing all affect this genotypic diversity.

Selective pressures can include biotic and abiotic stresses, the prevalence of which can vary at different times of year and from year to year. These pressure can select for favourable alleles in a mixed population or favour new advantageous mutations over the standing genetic variation. Grime (1977) defined classic strategies for plant survival in selective environments including competition (C), stress tolerant (S) and tolerant to ruderal/frequently-disturbed environments (R) the CSR model. Plants that are competitive tend to have rapid growth with extensive lateral spread above and below ground and low seed yield. Stress tolerant plants tend to have slow growth with small leaves and low seed yields. Ruderal plants tend to have rapid growth, with small stature and large yield. Most Brachypodium accessions seem to favour a ruderal strategy, however the diversity of life strategies, growth rates, and plant architectural traits seen suggest that competition and stress tolerance are also important selective pressures.

Isolation by distance can cause populations to diverge over time and has a major impact on Brachypodium diversity. This differentiation may occur via physical barriers, such as a mountain range, ice sheet or seas. This can be seen in the East West divide within each *B. distachyon* subspecies. However, isolation by distance may also occur from a difference in reproductive timing as different life strategies initiate flowering at different times of year, eliminating the chance of cross-pollination. This could be a factor in the division of the A and B subspecies in *B. distachyon*. Post reproductive barriers could also be involved, such as genetic incompatibilities and hybrid breakdown.

The timing of germination and reproduction, also know as life strategy, are good examples of phenological variation that are strongly shaped by natural selection in local climates. Brachypodium accessions collected from high altitude and high latitude sites are generally strong ‘vernalisation requiring’ ecotypes (Schwartz et al. 2010). This allows extended vegetative growth and the synchronisation of flowering with the more favourable conditions of late spring rather than risking damage to cold-sensitive floral structures in winter. In Mediterranean regions, winter rainfall determines the growing season for ‘rapid cycling’ ecotypes to avoid the hot dry summers. Accessions from hot arid climates with intermittent rainfall as well as disturbed agricultural site also tend to harbour ‘rapid cycling’ ecotypes with fast generation times, low dormancy and low vernalisation requirements, e.g. Bd21-1 (Barrero et al. 2012). This fast generation time allows a full life cycle to occur when moisture is available, possibly two times a year, while enduring dry periods as seed. Though rare, Brachypodium accessions from warmer, wet summer regions are later flowering allowing more vegetative growth and increased summer seed set. Some natural accessions may also have the ability to produce different phenotypes in different environmental conditions because of phenotypic plasticity. For example, accessions from variable climates may have some vernalisation requirements, but also a strong photoperiod response such that if there is a cold winter, flowering can be promoted, but in a mild winter flowering is initiated by a strong photoperiod response.

#### Case Study: Flowering Time

An experiment to investigate the genotypic and environmental basis for the diversity of flowering time of *B. distachyon* was undertaken at ANU in 2014. Plants were grown in specially modified growth chambers to simulate a temperate region with a cold winter and hot summer. Chambers were set for either a Winter or Spring germination. The modified chamber controls allowed us to program down to one minute changes in light intensity, light spectrum, temperature and humidity; mimicking diurnal and seasonal changes in climate. In this experiment 256 accessions were grown in each chamber. Ear emergence was chosen as an indicator of transition to reproduction as many accessions flowered within the ear and hence were difficult to score for flowering. The Winter germination chamber had cold (5°C nights) resulted in the vernalisation requirements of all lines being met and the majority of lines reaching ear emergence between 100-160 days post emergence (Figure 3a). However, in the Spring germination treatment the transition to reproduction was more variable. The majority of lines in the Spring germination reached ear emergence fairly quickly between 60 - 120 days post emergence. However, although there were still a few weeks with cold nights, the accessions requiring stronger vernalisation did not reach ear emergence by the end of the experiment at 180 days (e.g. Tek1, ABR4, UKR-99-137). Interestingly, while the life strategy of the majority of lines remained fairly unchanged between environments (early flowering), those accessions from Western Europe tended to show more phenotypic plasticity between environments with a wider range of delay in ear emergence with the Winter sowing than the Spring sowing.

**Fig. 3.**
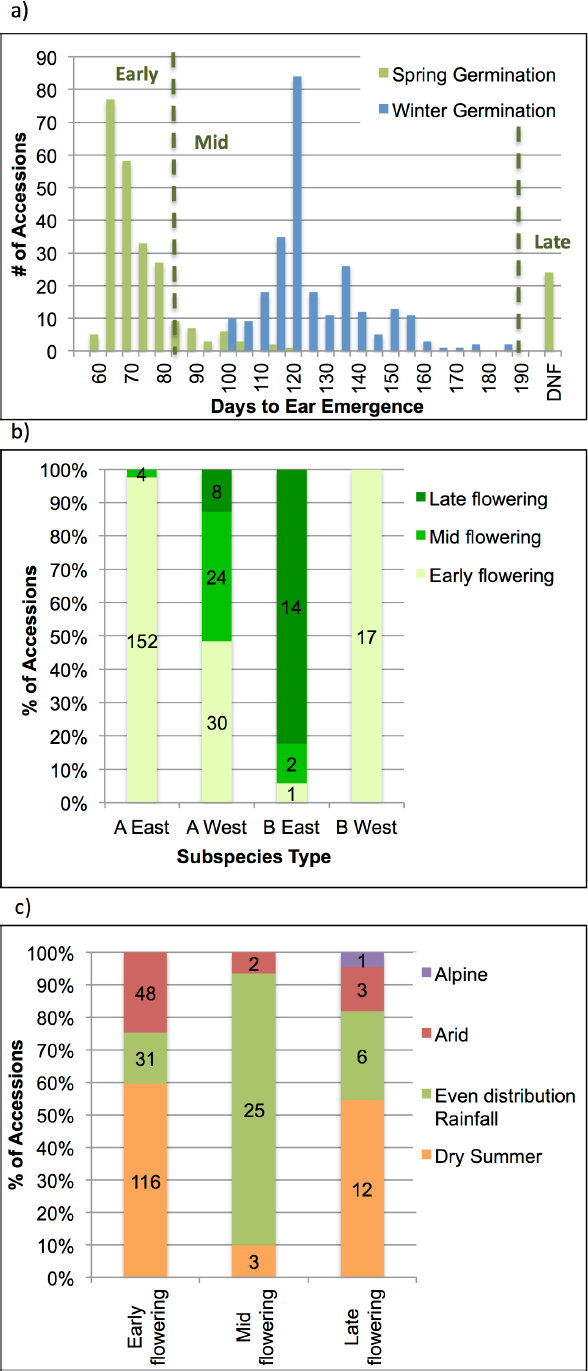
Life strategy of *B. distachyon* accessions is associated with subspecies, region and climate type at the collection site. 256 Brachypodium accessions were grown in simulated growth conditions to mimic a Winter and Spring sowing date. Days from seedling emergence to ear emergence was monitored as an indicator of transition to the reproductive stage. a) Comparison of life strategy in the two environments; dashed lines indicate the cutoffs for classification of accessions as early, mid or late flowering based on their flowering time under the Spring germination. b) Subspecies and geographic classification of accession show varying distributions of early, mid and late flowering accessions; numbers in columns indicate number of accessions in each category and c) Early, middle and late flowering groups of accessions are sourced from different proportions of climate types.

The accessions were then classified as early, mid or later flowering based on the days to ear emergence in the Spring sowing (Figure 3a). When these groupings are compared to the subspecies classifications we can see there is no firm trend for flowering time between the subspecies, as the patterns differ significantly in the Eastern European and Western European subsets (Figure 3b). For example, while accessions from the A subspecies tend to be early flowering in Eastern Europe, a mix of life strategies are present in Western Europe. This mix would indicate that different flowering time alleles have evolved in the different subspecies. Making crosses between accessions from different geographic regions and also between subspecies within a region would be needed to genetically map divergent flowering time alleles. Fortunately multiple RIL sets are available spanning this natural divide. Just as flowering time is variable within subspecies and geographic regions, it also varies with climatic type at the site of collection. When we categorise our accessions by climatic region, the life strategy patterns are not fixed based one on selective pressures in these climates alone. Accessions from climates with dry summers can be early or late flowering (Figure 3c). A similar trend is seen for accessions from arid regions, where a wide range of flowering times are seen. Mid flowering accessions tend to be from climates with rainfall evenly distributed through the year, indicating that the accessions with long growing seasons and no vernalisation requirements have a fitness advantage in this climate type. An alpine accessions was late flowering, suggesting that vernalisation is needed for survival in this environments. We must keep in mind sampling bias in any collection. The observed trends can be influenced by when and where the collections are made. Netherless, there are trends with exceptions, illustrating the impact of the genotype and climate on the geographic distribution and phenotypic expression of accessions.

Understanding fitness of an individual accession in a particular climate is complex, complicating studies aimed at elucidating the genetic architecture of adaptive traits. Luckily, new approaches have been developed that control for genetic and spatial structure, which can result in outcomes that have applications in crops species breeding programs, weed management and ecosystem restoration. Next we outline two of these approaches: Genome Wide Association Studies (GWAS) and Landscape Genomics.

### 2 Using Local Adaptation to Identify Causative Alleles

#### Genome Wide Association Studies

Natural populations provide a valuable resource in the diversity of genotypic variation that has been found in extreme environments. Historically these populations have been studied for phenotypic diversity in small numbers of accessions using SSR and microsatellites. The development of comprehensive next generation sequencing has meant that thousands of accessions can now be screened and powerful subsets selected for Genome Wide Association Studies.

Genome wide association studies (GWAS) were originally developed for studies of human disease to find causative mutations (Klein et al. 2005). The aim of GWAS is to identify the genetic architecture of phenotypic traits of interest in a certain population. This can include the number of alleles associated with the trait, whether one allele affects multiple traits and the strength of individual alleles in accounting for the trait of interest. By comparing different environments we can also identify which alleles are important in each environment, which are robust across environments and which allow for plasticity of the trait between environments. GWAS have been performed in a number of plant species including Arabidopsis, maize, rice, barley, and sorghum (e.g. Atwell et al. 2010; Wen et al. 2014; Huang et al. 2010; Pasam et al. 2012; Morris et al. 2013). These studies have successfully identified causative QTLs for many traits including disease resistance, proline accumulation, flowering time, starch profile, plant height and grain weight.

##### Genotype Data

To perform GWAS you must have genotyped a large number of SNPs tagging the many haplotype blocks that are differentially assorted in the sample being studied. The density of SNPs needed depends on ancestral recombination in the genome and will affect the resolution of the QTLs identified. To get high density SNP genotyping whole genome re-sequencing can be used by randomly shearing DNA and ligating barcoded adaptors. Like GBS, samples are then multiplexed and run on an Illumina HiSeq platform. Low pass sequencing (∼1-2X) is sufficient as missing genotypes at common SNPs can be imputed based on linkage disequilibrium (Halperin and Stephan, 2009). GWAS must also be performed on a sizable number of accessions, at least 100 but optimally ∼300, to maximise statistical power. SNPs are usually filtered at 5% minor allele frequency. A genome scan is then performed testing each SNP for association with the quantitative trait while controlling for ancestral relatedness among samples. Once a QTL are identified to perhaps 100kb resolution, candidate genes can be selected. Ultimately the causative alleles can be confirmed by transgenic complementation of knockout lines with different functional alleles.

GWAS has some advantages over traditional bi-parental mapping (e.g. RILs), nested association mapping (NAM) or multi-parent advanced generation intercrossed (MAGIC) populations in that the source of genetic variation is much greater as its not just limited up to a couple dozen founder accessions, but comes from hundreds. Furthermore, the resolution of mapping of the causative allele can be much greater due to the higher frequency of recombination events in a large ancestral population evolved over centuries compared to a hand-crossed population with only a handful of generations of crossing. However, rare alleles of large effect contribution noise for GWAS. In addition, population structure from non-random mating can cause false-positives at loci associated with the causative one. Even when appropriately controlled for statistically, population structure can cause false-negative results by downplaying the effect at the truly causative locus.

There are a couple approaches to mitigate the effects of population structure. The first is to balance the structure. GBS can be performed on a very large number of diverse lines from across the world. Once the structure of the global population is known, a core GWAS set can be selected balancing membership roughly equally from across different lineages (Figure 4a). Whole genome sequencing is performed on this GWAS set, to increase resolution of mapping causative alleles. A kinship matrix is also made from the SNP data to include in the statistical analysis to reduce the rate of false positives. The second approach is to perform the same initial GBS screen of a wide range of accessions, but then to select a region within a population structure group or a hybrid zone where the groups have become well mixed. The resulting population sample be less confounded such that segregation at the causative alleles will be largely uncoupled from the background genetic variation. These approaches are further described in Brachi et al (2011) and an example of its success in Arabidopsis is given by (Li et al. 2010: Baxter et al. 2010; Horton et al. 2012).

**Fig. 4.**
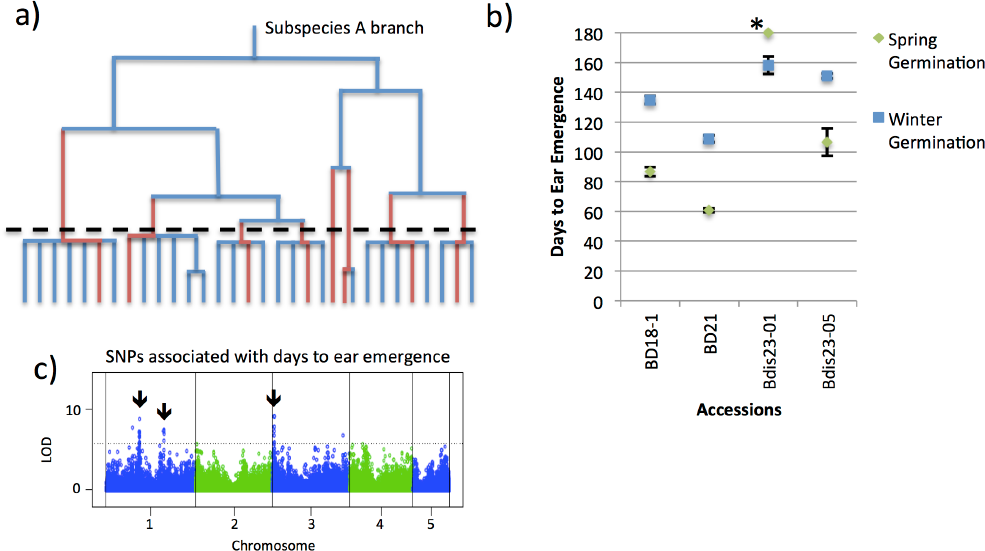
Considerations when performing GWAS in Brachypodium. a) Selection of a balanced set of accessions is key for successful GWAS. Initial GBS of all available accessions facilitates selection of a subset of lines for intense phenotyping and whole genome sequencing b) Replication of a few key lines allow for the heritability of a phenotypic trait to be estimated. Traits with large environmental variance will return poor GWAS results. Points are the average time to ear emergence of 3-6 biological replicates ± S.D.; * did not flower in the Spring germination c) A genome wide screen of SNPs identifies those associated with a quantitative trait. Manhattan plot with arrows showing SNPs associated with days to ear emergence with a Spring germination. A balanced set of 95 accessions from the A subspecies was used in analysis. The dotted line indicates the 5% empirical genome-wide significance threshold.

In consideration to the first approach outlined above, GWAS cannot be performed across the A and B subspecies due to a high level of genome level differentiation. Hence we suggest that GWAS sets are developed for each subspecies separately. Furthermore, the current *B. distachyon* public collections contain a lot of groups of very closely related accessions, e.g. BdTR1_ lines. Selection of a GWAS set must also limit family structure. Including closely related accessions limits recombination and introduces bias towards the common alleles in that family. The second approach of finding a well-admixed population may be challenging due to the selfing nature of Brachypodium. However, it is our experience that several field sites and even some maternal lines are segregating both genotypes and phenotypes, indicating that this strategy is a possibility.

##### Phenotyping Data

There are a number of GWAS requirements for phenotype data. The traits must be due to common genetic variation rather than multiple rare variants. It is helpful to measure heritability, or the proportion of total phenotypic variation (Vp) due to genetic variation (Vg) among lines. A simple way to estimate heritability (H2) is to subtract environmental variation (Ve) within replicate inbred lines, H2 = Vg/Vp = (Vp-Ve)/Vp. If the variability within a genotype is low then more of the variability seen in the phenotype will be due to genetic factors (Figure 4b). Alternatively the pseudo-heritability is the amount of phenotypic variation explained by the kinship or pairwise genetic diversity among lines and can be calculated prior to GWAS.

Due to the large number of accessions to be phenotyped for GWAS, the phenotypes must also be quantifiable in a high throughput manner. For some measures, such as plant height or ear emergence this can be done manually with relative ease. However, for traits that require continuous measurement, such as growth rates, or those where measurements and analysis are very time consuming, such as photosynthetic measures, high throughput phenotyping systems can be very beneficial. One such system is the PlantScreen phenotyping platform (Brown et al, 2014). This platform allows up to 300 plants to be screened at a time; including individual pot weighing and watering, thermal imaging, RGB stereo imaging and chlorophyll fluorescence. This system allows us to monitor abiotic stress tolerance of accessions by measuring photosynthetic efficiency and photoinhibition; water use; stomatal conductance/transpiration cooling; and photo-protective mechanisms such as pigment accumulation and non-photochemical quenching (NPQ).

##### Genotype by Environment Interactions

An important extension of GWAS is to understand how the growth environment affects the genetic architecture of traits. Certain loci may be more important in the expression of a trait of interest in particular environments. A simple example is where a vernalisation-sensitive allele may be important in controlling flowering time in an alpine climate, but a photoperiod-sensitive allele may be more important in a lowland climate. The interaction between genetic loci may also vary between environments. Understanding the genetic architecture of phenotypic plasticity is also of great interest, especially in a world where climate variability and change is accelerating. Hence it is important to consider how the growth environment will influence the phenotypes seen in GWAS experiments and how this will relate to extrapolation of results to applications of plant breeding or ecological work.

Using dynamic growth chambers, such as the SpectralPhenoClimatron (Brown et al, 2014; Li et al, 2006), in combination with climate modelling software such as SolarCalc, we can undertake GWAS to simultaneously compare contrasts such as life strategies at different times of year (Figure 3, Li et al, 2010); growth and yield in current and future climates (e.g. Li et al, 2014); and the effect of fluctuating light, as would be seen under a forest canopy, on growth and initiation of photoprotective mechanisms.

##### Applications

GWAS allows us to understand the genetic architecture of interesting phenotypic traits, whether it is flowering time, biomass production, components crop yield or abiotic stress tolerance. With this approach we can identify segregating alleles in natural populations causing advantageous phenotypes (Figure 4c). This understanding can then be applied to breeding better crops by selecting genotypes with predicted to have optimal phenotypes, aka crop design (Huang and Han, 2014). Understanding how the genetic architecture of traits is affected by environment allows us to select accessions with the appropriate genetic makeup to thrive in specific environments, current or future. This can apply to both agriculture and foundation species of natural systems as climate change is occurring more rapidly than traditional breeding by phenotypic selection keep up with (Hoffmann et al, 2015).

In summary, GWAS is a powerful method to dissect the genetic basis of complex traits and the dependence on the growth environment. It is of interest to know if these are adaptive traits and contribute to yield and fitness in particular environments as this can help predict invasive potential and resilience under environmental change. Genes controlling adaptive traits may show a signature of selection on the landscape if one allele provided an advantage in a particular environment and/or the other allele was detrimental in an opposing environment. Indeed direct associations between environmental variables at geographic locations and alleles at quantitative trait loci have been found, confirming their role in adaptation (Li et al, 2010 and others). This opens the door to a genome scan to naively identify adaptive loci associated with environmental variables. This could work without knowledge of what traits are under selection. Though promising, this so called Landscape Genomics approach, has caveats including needing to control for background variation due to population structure. Geographic location is confounded with many environmental variables limiting our ability to separate selection at adaptive loci from isolation by distance that shapes the entire genome.

#### Landscape Genomics

Landscape Genomics is a growing field in plant biology integrating the sciences of geographic and climate mapping with population genomics and association studies. Teams of researchers in this field must have a diverse set of skills ranging from the lab to field studies uniting theory and computational biology. They must collect and analyse large data sets to correlate spatial and environmental variables with perhaps millions of genetic loci while accounting for genetic relatedness and demographic processes affecting the whole genome. Landscape genomics combines high resolution spatial/temporal climate data from the sites of collection with second generation sequencing technology to 1) explore the differences in climatic niche breadth among genotypes; to 2) select locations with variable microclimates and well mixed genetic variation; and 3) to associate particular climatic variables (e.g. winter temperature or precipitation levels) with specific adaptive alleles. Landscape genomics can be applied to almost any species in the natural environment, including animals, fungi, and plants. Some of the most successful studies are often of model species and their close relatives, which usually have annotated reference genomes and large developed collections that are already sequenced (Hancock et al, 2011, Fournier-Level et al, 2011).

Landscapes are longstanding experiments of natural and, increasingly, artificial selection. Landscape genomics dissects and describes the underlying adaptive genetic loci through association with a suite of environmental variables describing the microclimate within a species range. Initially, populations are founded with limited genetic diversity. New mutations are initially rare and most disappear due to genetic drift. When new genetic variation arrives via migration, it brings a genome’s worth of standing genetic variation. Even in largely selfing species, outcrossing will eventually happen and new allelic-combinations will be released for selection across the environment. As in synthetic crosses, the recombinant chromosomes initially have extended haplotypes from the founders that are ultimately broken down through subsequent generations and further outcrossing. There may now be opportunity for local range expansion as the diverse population spreads into new microclimate, soil and/or habitat types. As the ecological space fills, competition will favour plants with adaptive alleles in their most suitable habitat. Neutral genetic variation will have a largely random pattern across the landscape. Across the genome, a trend of isolation by distance may develop if gene flow is limited. Ideally this would be in a direction orthogonal to the environmental gradient. At this stage a genome scan for SNP alleles in association with local environment can identify adaptive loci. The allele effect at a locus multiplied by the strength of selection determines the ability to identify and adaptive QTL. The size of the adaptive locus is determined by the amount of recombination, which should narrow as the population ages. Control of false positives due to population structure can be performed by statistically accounting for overall relatedness across the genome in a manner analogous to GWAS. Across large geographic regions multiple rare variants explain adaptation limiting Landscape Genomics in much the same way as GWAS.

One advantage of annual plants is selection acts quickly at each generation. It can be observed at a locus when a few ancestral haplotypes are competing and gene flow mixes the background genetic variation. By understanding the genetic basis of local adaptation, we may finding allelic variants that allow some groups to be specialists, while other alleles may be important for generalists (Storz, 2005). Brachypodium has a large geographic range, mostly spanning Mediterranean-like climates in Europe, North Africa and Eurasia. It is similar to *Arabidopsis thaliana*, which has genetic diversity associated with geographic and climatic space (Platt et al. 2010; Horton et al. 2012; Banta et al. 2012). Each subspecies of Brachypodium distachyon would be expected to show similar associations with adaptive alleles across geographic and climatic space.

So the next step is identifying which populations and climate regions are suitable for Landscape Genomics. Different genetic lineages may occupy different geographic ranges and corresponding climate envelopes. These can highlight potential adaptive differences however alleles providing the advantage may be fixed within each lineage. Hybrid zones between groups or long range dispersal followed by admixture could provide the opportunity to uncouple adaptive alleles from their genomic background. Digital herbarium records are a great place to start describing the climate range of a species utilizing existing collections. Records often have meta data about microclimate, some phenotype data including whether the plant was flowering, in addition to when and where the plant was collected. This can aid a researcher’s decision about when to travel for collecting, what traits to look for and what locations they occupy. With collection points, researchers can also travel back to the same location or similar locations based on model predictions like those created in computer programs to predict niche breadth using climate envelopes (Phillips, 2006; Joost, 2007; Banta, 2012).

#### Determining the geographic range and climatic niche breadth of divergent genetic lineages

While the geographic range and environmental tolerance of a species can be wide, the breadth of environmental conditions a subspecies, or major genetic lineage, can inhabit may be relatively small. Understanding climatic breadth within species can inform growers about what crop varieties are suitable for their locations and can also aid seed collectors about new locations for future collections. Maximum Entropy (MaxEnt) software is a common research tool for inferring and predicting species distributions and tolerances of environmental factors (Phillips, 2006; Phillips, 2008). The inputs are flexible and can be manipulated by the user. Data is often sourced from WorldClim, a global grid of climate layers (WorldClim, 2015). A subset of biologically relevant variables, BioClim, are often used in ecological studies including species range modeling programs (BioClim, 2015). These variables represent growth limiting factors such as temperature and precipitation and their seasonality. MaxEnt couples environmental data from training locations as inputs and compares these realised niche variables with all locations outputting geographical heat maps indicating where similar ecological niche exist. The results describe a species fundamental niche, which is the climate limited range where a species might be able to persist. MaxEnt outputs response curves of climate variables and uses permutations to rank the relative contribution of each variable when determining the niche. The program uses the model to predict sites left out of the training data. This cross validation also allows comparisons between models. Different models can be made and compared which incorporate population genetic information. This is done by defining different sets of training locations based on the occurrence of particular genetic lineages. Figure 5a shows the geographic range of the Australian Brachypodium fundamental niche generated by MaxEnt trained with the Bioclim climate variables at 80 locations that comprise our current collections. Mean Temperature of Wettest Quarter (WorldClim 8) was the strongest factor determining the distribution (BioClim, 2015).

**Fig. 5.**
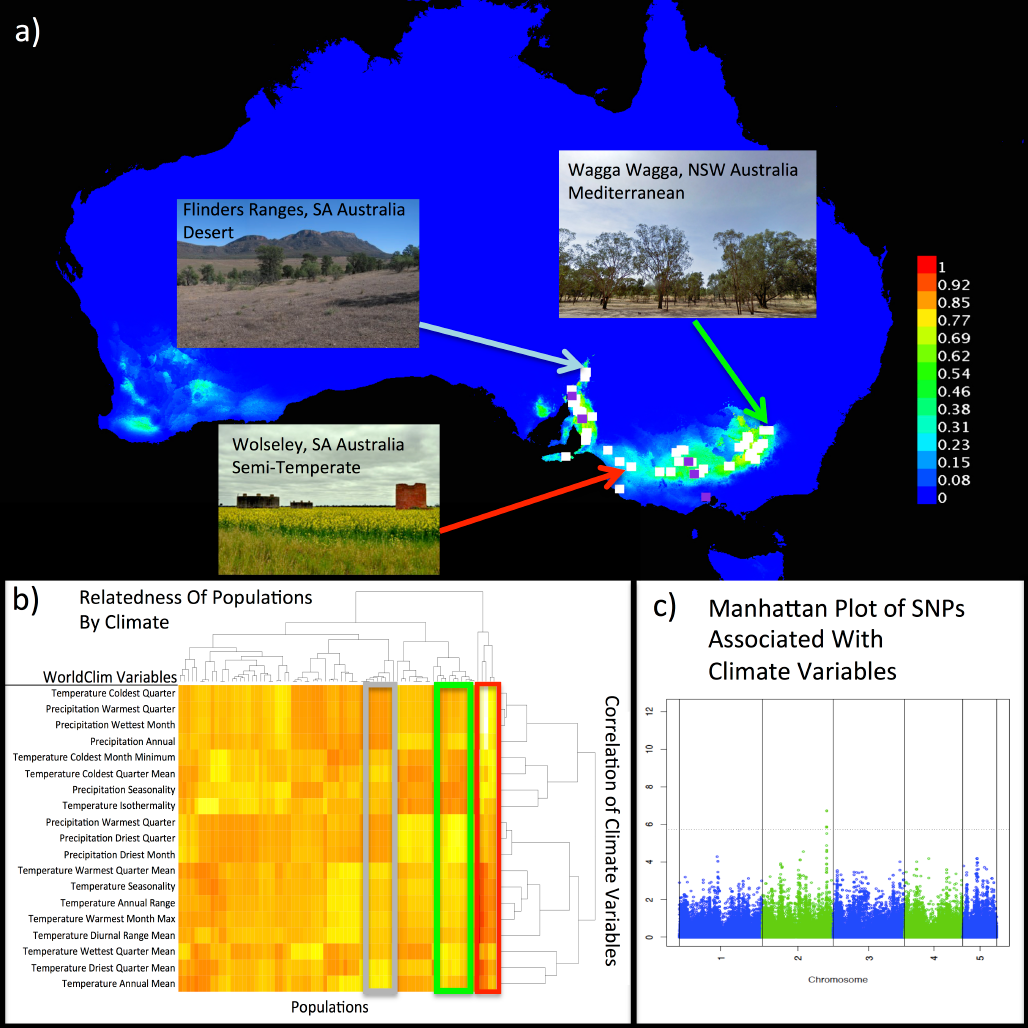
Predicting species distribution and detecting segregating SNPs by climate input. a) MaxEnt produced probable distribution of Brachypodium in Australia using 80 geographic locations as presence only data correlated with WorldClim/BioClim variables. b) 80 Australian populations collected by the Borevitz lab of *B. distachyon* complex to show the environmental relatedness of each population (no genetic data used). The vertical dendrogram describes environmental variable correlation across all populations. The horizontal dendrogram describes individual population clustering by their environmental relatedness to inform researchers about where certain genotypes/haplotypes might grow. If specific haplotypes are found in multiple locations one can dissect which strains are more likely to be pervasive. Likewise the variation and possible selection pressure one or more environmental variable(s) may have across many populations. Using a custom R script, we normalize 19 WorldClim environmental variables point sampled across locations. Variables with the smallest normalised distribution from the mean are likely to be a strong selection pressure species-wide. Variables with larger distribution could be random, but also could be associated with specific alleles. c) A cartoon representation of SNPs associated with environmental variables, calculated much the same way as a LOD with phenotype data.

##### How do neutral forces affect the genetic signal of climate adaptation?

The climatic niche breadth of a genetic lineage is the result of both adaptive and neutral forces. With limited migration, founder effects and isolation by distance, prevent our ability to separate the chance historical demographic effects on the entire genome from the adaptive genes. We must consider the genetic divergence and populations structure when selecting locations for intensive sampling. Recently founded populations may not have enough diversity to identify haplotype blocks associated with climate variables while diverged and structured populations may not allow adaptive loci to be partitioned from background variation. Geographic distance can often explain genomic isolation (eg Mantel test) within a species and can be the major factor at large, intercontinental scales leaving little ability to identify adaptive loci. Mantel tests can be performed at regional scales across clusters of sites to identify groups where isolation by distance is not a dominant factor. Partial mantel tests can be used to further account for genomic isolation by environment in addition to geographic distance. These background effects are essential to control for when looking to associate particular adaptive genetic loci with specific environmental variables.

A more advanced approach than Mantel tests takes advantage of the software package Popgraph in R. It is a valuable tool for describing genetic relatedness between populations and environmental layers. The software works similarly to the PCoA algorithm, only it keeps the collection locations static on a map and uses visual tools to show genomic relatedness across geographic locations (Garrick et al, 2013). MaxEnt, Mantel tests, Popgraph, and other methods can provide comparisons of environments, geographic space, and genomic variation.

##### Identifying which climate variables

When overall climate is found to be associated with genomic variation, it is of interest to investigate which particular environmental variables are the drivers of natural selection. This can be done by clustering environmental variables across geographic locations. Figure 5b shows an ordered heat-map of 80 collections locations by 19 WorldClim environmental variables. The climate data is provided via Atlas of Living Australia (ALA) for points at sample locations. In this heatmap there are two large clades of environment types with 5 sub-groups. While there is similarity among environments at a local scale extending East and West, there is large split North to South. Three geographic regions are highlighted which represent three major climate types. The Mediterranean regions (green) cluster in climate space near desert regions (grey) while temperate regions (red) are separated. This is despite the fact that Mediterranean regions are geographically separated from desert regions.

##### Testing adaptation to climatic range

Typically common gardens and reciprocal transplant studies are used to test for local adaptation where the home genotype performs superior to the genotype from farther away (e.g. Clausen et al. 1940). Provenance trials do this on a larger scale, evaluating phenotypes from a broader range of genotypes across many locations. These studies are massive and are certainly hampered by starting conditions and weather variation. As described above, one solution is to use growth chambers that can mimic natural climatic conditions, without weather noise. The SpectralPhenoClimatron, allow fitness traits to be measured in multiple target environments and using high-throughput phenotyping techniques.

##### Scanning for Adaptive Genetic loci controlling survival or fitness

Measuring fitness directly is difficult. It is the result of selection at many different life stages and often differs across environments and variation at many loci can contribute. An indirect way is to use the presence of a plant in a location as an observation that a particular combination of alleles can survive there. When a large collection of plants spanning a suitable range of climate variables are sequenced, one can use environment data as a fitness phenotype to and test if certain alleles associated. A genome wide association study scan can then be performed with climate data to identify adaptive loci. Figure 5c shows a cartoon example of specific loci associating with Precipitation in the Warmest Quarter.

#### Future Collections

Extensive public collections exist, however many geographic regions and climate niche remain unrepresented or undersampled. Having those spaces filled would be beneficial for the whole Brachypodium community. As mentioned above there is a strong division in the current public collection between Eastern and Western European accessions. Hence, it would be advantageous to have collections across the Middle East and North Africa as it is highly possible that *B. distachyon* complex species were pushed south during the last ice age and extant lineages could be sources of new maternal lines for research. Central Southern Europe might provide a source of admixture populations between the Eastern and Western genotypes or completely new genotype groups. With genotype data, we can focus further collections on hybrid zones and polymorphic sites to increase the number of recombinant genotypes. These genotypes provide unique opportunities to study natural selection on adaptive traits. They provide natural genetic mapping resources to dissect complex adaptive traits. Admixed populations also inform the evolutionary history of Brachypodium and provide an opportunity to test natural selection on segregating variation.

To study the natural variation of stress tolerance in *B. distachyon*, it would be good to have further collections from more extreme environments. This would include higher altitude/latitude environments such as Northern Europe and more arid environments in the Middle East and North Africa. Areas with multiple, recent introductions are also of interest as they may provide examples of strong selection on standing variation.

## Conclusion

Brachypodium has a large genetic diversity with its distribution spanning a wide range of climatic regions. Through Landscape Genomics we can use this diversity to predict the niche breadth and potential range size of a particular ecotype. We can also start to understand the genetic basis of fitness/adaptation to this niche. Genome Wide Association Studies utilises Brachypodium genetic diversity in combination with its broad phenotypic diversity to elucidate the genetic architecture of traits of interest and how this architecture varies in different climate types. Both techniques rely on a thorough understanding of the population structure of available collections with particular attention to sub-genomic divisions. Future developments in these techniques will rely heavily on the expansion of the collection of publicly available accessions and development of bioinformatic and statistical methods to increase the power and accuracy of both the Landscape Genomics and GWAS approaches. However, via recently developed techniques, the use of these approaches in Brachypodium has much potential for applications in crop design and ecological restoration projects for future climates.

## Acknowledgments

We would like to acknowledge our collaborators: Location data and seed - Shuangshuang Liu of the Kent Bradford lab at UC Davis United states, Pilar Catalan from the University of Zaragoza in Huesca Spain, Luis Mur at Aberystwyth University at Aberystwyth Wales, Dave Garvin from University of Minnesota/USDA United States and John Vogel from JGI/UC Berkeley.

## Abbreviations

GBS: genotyping by sequencing
GWAS: genome wide association studies
QTL: quantitative trait loci
MaxEnt: Maximum Entropy
SNPs: single nucleotide polymorphisms

## Glossary

Accession: A collection of seeds from one location. This includes bulk collections and maternal descent lines
Ecotype: An individual or group whose genetic distinction is strongly associated to an environment or type
Genotype: This general term is used either to describe the genotype at a locus such as a SNP (AA, Aa, aa) or a background whole genome genotype which can have levels of species, subspecies, population genetic structure group, family, individual maternal line)
Phenotype (qualitative and quantitative): Measurable traits expressed by plants
Population: Non-random mating between groups within a specified geographic space
Subspecies: In this paper, subspecies is a major hierarchical cluster of genotype groups and their respective families and/or genotypes. Subspecies could interbreed but don’t in natural environment due to some sort of natural barrier

## Footnotes

**More details can be found here:** http://borevitzlab.anu.edu.au/resources/populations/

